# Structural basis of human IL-18 sequestration by the decoy receptor IL-18 binding protein (IL-18BP) in inflammation and tumor immunity

**DOI:** 10.1101/2022.02.10.479912

**Authors:** Sammy Detry, Julie Andries, Yehudi Bloch, Cem Gabay, Danielle Clancy, Savvas N. Savvides

## Abstract

Human Interleukin-18 (IL-18) is an omnipresent pro-inflammatory cytokine of the IL-1 family with central roles in autoimmune and inflammatory diseases, and serving as a staple biomarker in the evaluation of inflammation in physiology and disease, including the inflammatory phase in COVID-19. The sequestration of IL-18 by its soluble decoy receptor IL-18 Binding Protein (IL-18BP) is critical to the regulation of IL-18 activity. Since an imbalance in expression and circulating levels of IL-18 is associated with disease, structural insights into how IL-18BP outcompetes binding of IL-18 by its cognate cell-surface receptors would be highly desirable. However, the structure of human IL-18BP in complex with IL-18 had remained elusive. Here, we elucidate the sequestration mechanism of human IL-18 mediated by IL-18BP based on the crystal structure of the IL-18:IL-18BP complex. These detailed structural snapshots reveal the interaction landscape leading to the ultra-high affinity of IL-18BP towards IL-18 and identify substantial differences with respect to previously characterized complexes of IL-18 with IL-18BP of viral origin. Furthermore, our structure captured a fortuitous higher-order assembly between IL-18 and IL-18BP coordinated by a disulfide-bond distal to the binding surface connecting IL-18 and IL-18BP molecule from different complexes, resulting in a novel complex with 2:2 stoichiometry. This tetrapartite assembly was found to restrain IL-18 activity more effectively than the canonical 1:1 complex. Collectively, our findings will provide a framework for innovative structure-driven therapeutic strategies and further functional interrogation of IL-18 in physiology and disease.

**Significance statement:** Elevated levels of interleukin-18 (IL-18) have long been implicated in numerous inflammatory diseases while also displaying potent anti-tumoral activities. Recent research on COVID-19 has now underscored the role of IL-18 and IL-18 binding protein (IL-18BP), a soluble receptor serving to regulate IL-18 activity, as key players in viral immunity and as promising biomarkers and predictors of disease severity. In this work, we present detailed structural insights into how human IL-18 and IL-18BP interact thereby completing the structural repertoire of IL-18 in complex with its cognate human receptors and viral decoy receptors. Our findings will support structure-based efforts to either disrupt or enhance the interactions of IL-18 with its cognate receptors for therapeutic purposes.

## Introduction

Originally discovered in mice as Interferon-γ-inducing factor (IGIF), Interleukin-18 (IL-18) was found to be a potent inducer of interferon-γ (IFN-γ) production in synergy with IL-12 upon exposure to intracellular pathogens (1–3). It was subsequently cloned and identified as an 18 kDa cytokine belonging to the IL-1 cytokine family, and renamed to IL-18 (4). Similar to other IL-1 family cytokines, IL-18 is expressed as an inactive precursor protein that requires cleavage by caspase-1 for full biological activity (5, 6). While approximately 80% of pro-IL-18 is retained intracellularly, the rest is released by macrophages/monocytes and dendritic cells after maturation by caspase-1 (7, 8). Mature IL-18 initiates signaling by establishing a tripartite complex with IL-18 receptor alpha (IL-18Rα) and IL-18 receptor beta (IL-18Rβ), the latter also known as IL-18 receptor accessory protein and IL-1R7. The heterodimerization of the receptors’ intracellular Toll/IL-1 receptor (TIR) domains triggers recruitment of the adaptor protein myeloid differentiation primary response 88 (MyD88) which further activates the downstream signaling cascade through IRAK-1/4, TRAF6 and NFκB (9–12). IL-18 is mainly involved in the activation of NK and T helper 1 (Th1) cells and IFN-γ production in response to intracellular bacteria or viruses (13, 14).

Due to its highly proinflammatory and pleiotropic activity, IL-18 signaling is highly regulated at many levels to prevent uncontrolled inflammation. Akin to other IL-1 family cytokines, this regulation is achieved through gene regulation (15), caspase-1 activation (5, 6) and importantly, is also mediated by a soluble decoy receptor (16). Once IL-18 is activated and released from immune cells, its availability is mainly regulated by sequestration by IL-18 binding protein (IL-18BP), a dedicated soluble decoy receptor-like protein capable of blocking the biological activity of IL-18 (17). Even though IL-18BP functions as a decoy receptor, it is not homologous to any extracellular domain of the IL-18 receptors and is encoded as a separate gene (18), an oddity in the cytokine field (19). However, bioinformatic approaches have proposed that IL-18BP is evolutionary related to TIGIRR-1 (aka IL-1R9) and is thus regarded as a member of the IL-1 receptor family. In addition, sequence and functional homology lead to the identification of IL-18BP orthologs in all orthopoxviruses (20, 21), including ectromelia virus (ectvIL-18BP) and Yaba-Like Disease Virus (yldvIL-18BP) (22, 23), that act as virulence factors by attenuating immune responses mediated by IL-18.

Not only is human IL-18BP present in the serum at a 20-fold molar excess (24), it also has an exceptionally high affinity (K_D_∼300 pM) for IL-18 (25), in contrast to IL-18Rα’s affinity towards hIL-18 which is reported to be several orders of magnitude weaker (K_D_∼69 nM) (26). Intriguingly, the affinity of viral IL-18BPs for human IL-18 is considerably lower than human IL-18BP (K_D_∼1 nM), although they display picomolar affinity towards mouse IL-18 (25). As a result, IL-18 is sequestered by IL-18BP under homeostatic conditions, thereby preventing it from signaling through its receptors and evoking unwanted pro-inflammatory responses. Interestingly, IL-18-induced IFN-γ upregulates IL-18BP expression, creating a negative feedback loop to dampen and resolve inflammation (27, 28). Several (auto)immune diseases have been associated with increased levels of IL-18 such as rheumatoid arthritis (RA) (29, 30), Crohn’s disease (CD) (31, 32), and systemic lupus erythematosus (SLE) (33–35). As IL-18BP has a high IL-18 sequestration capacity (35), the balance between IL-18/IL-18BP and the concentrations of free IL-18, instead of total IL-18, are more relevant to measure to evaluate inflammatory responses (36). Elevated levels of free IL-18 have been identified in hyperinflammatory diseases such as macrophage activation syndrome (MAS), and systemic juvenile idiopathic arthritis (28, 37). Furthermore, in mouse models of MAS, IL-18BP-deficient mice developed more severe disease manifestations, pointing to the crucial role of IL-18BP activity in this setting (28). In addition, elevated levels of IL-18 in blood and bronchoalveolar lavage fluid (BALF) from coronavirus patients have been shown to correlate with COVID-19 disease severity and worse clinical outcomes (38– 40).

Unsurprisingly, IL-18 neutralizing antibodies or recombinant IL-18BP have been successfully used to mitigate IL-18 related pathologies (41, 42). Clinical trials using recombinant hIL-18BP (Tadekinig α) for Adult-onset Still’s disease (AOSD), rheumatoid arthritis (RA) and plaque psoriasis show promising safety and indicative signs of efficacy in patients with AOSD (43, 44). On the other hand, administration of recombinant IL-18 in mouse tumor models was shown to elicit favorable anti-tumoral effects in synergy with chimeric antigen receptor T (CAR-T) and immune checkpoint inhibitors (45). Whereas such findings provided a strong rationale for the therapeutic potential of IL-18 in cancer (46), its efficacy has not lived up to expectations (47), the culprit being the concomitant increase in the serum concentration of IL-18BP by up to 100-fold in patients (46, 48).

To aid further advancements in the mechanistic interrogation of IL-18 sequestration by IL-18BP and the therapeutic targeting of hIL-18 and hIL-18BP in infectious diseases, inflammation and cancer, structural details of their complex and of interaction interface to high resolution would be opportune. Here, we present the crystal structure of human IL-18 in complex with human IL-18BP. This study shows that hIL-18BP binds hIL-18 at the same epitope as IL-18Rα and viral IL-18BPs using a large hydrophobic patch flanked by two tightly fitting hydrophobic pockets complemented by salt bridges. A crucial advance in our crystallization efforts was the elimination of the flexible N-terminus and minimization of heterogenous glycosylation patterns of hIL-18BP while retaining functional and biological activity. Moreover, we discovered a novel disulfide-linked interface resulting in an unexpected tetrameric assembly of hIL-18 and hIL-18BP. The structural model presented here is imperative to validate previous models, map key differences between human and viral orthologs of IL-18BP and allow advancements in the design of novel inhibitors or antagonists.

## Results

### Human IL-18BP can be truncated to a bioactive core structure with reduced glycosylation

To enable structural studies of the human IL-18BP:IL-18 complex by X-ray crystallography we initially considered clinical grade human IL-18BP (Tadekinig alfa) that was used in a phase 2 clinical trial to evaluate the safety and efficacy of IL-18BP in adult-onset Still’s disease (43). However, due to the extensive glycosylation of this protein and the need to create glycan-engineered protein amenable to structural studies by X-ray crystallography we produced full-length hIL-18BP in transiently transfected HEK293S MGAT1 -/- cells allowing for N-linked glycosylation as Man_5_GlcNAc_2_ glycans (49) and mature hIL-18 in *E. coli*. Using size-exclusion-chromatography (SEC) and in-line multi-angle laser light scattering (MALLS) we found that full-length hIL-18BP was heavily glycosylated with nearly 50% of its mass accounted for by glycans (**Figure 1A** and **Supplemental Table** 1), consistent with predictions (**Figure S1A**). Furthermore, the distribution of mass across the SEC-MALLS peak for full-length hIL-18BP, the diffuse electrophoretic mobility of the purified protein in SDS-PAGE (**Figure 1A&B**), and large shifts in mass upon treatment with endoglucanases and O-glycosidases suggested heterogeneous glycosylation. Based on our experience with producing secreted glycoproteins with N-linked glycosylation and similar protein families in the HEK293T and HEK293S MGAT1 -/- cell lines for structural biology (50–53), we reasoned that this most likely originates from O-glycans and possible differences in N- and O-glycosylation site occupancy. Initial crystallization trials using purified hIL-18BP_FL_:IL-18 complex containing glycan-shaved hIL-18BP_FL_ and displaying apparent stoichiometry of 1:1 as evaluated by SEC-MALLS, did not lead to candidate crystallization conditions even when employing hIL-18BP_FL_:IL-18 complex up to 30 mg/mL. Such apparent high solubility during crystallization trials was deemed to be consistent with the observed extensive and heterogeneous glycosylation of hIL-18BP_FL_ (**Figure 1A** and **Supplemental Table** 1). Therefore, we sought to produce alternative versions of hIL-18BP that would be more amenable to structural studies. Sequence alignment of hIL-18BP with homologous viral IL-18BPs, for which structural information is available (22, 23), suggested that the expected core beta-sandwich domain of hIL-18BP (residues 63-171) might be flanked by N- and C-terminal stretches (residues 31-62 and 170-194, respectively) bearing intrinsic disorder and the majority of the predicted O-glycosylation sites (**Figure S1A** and S1B).

**Figure 1.**
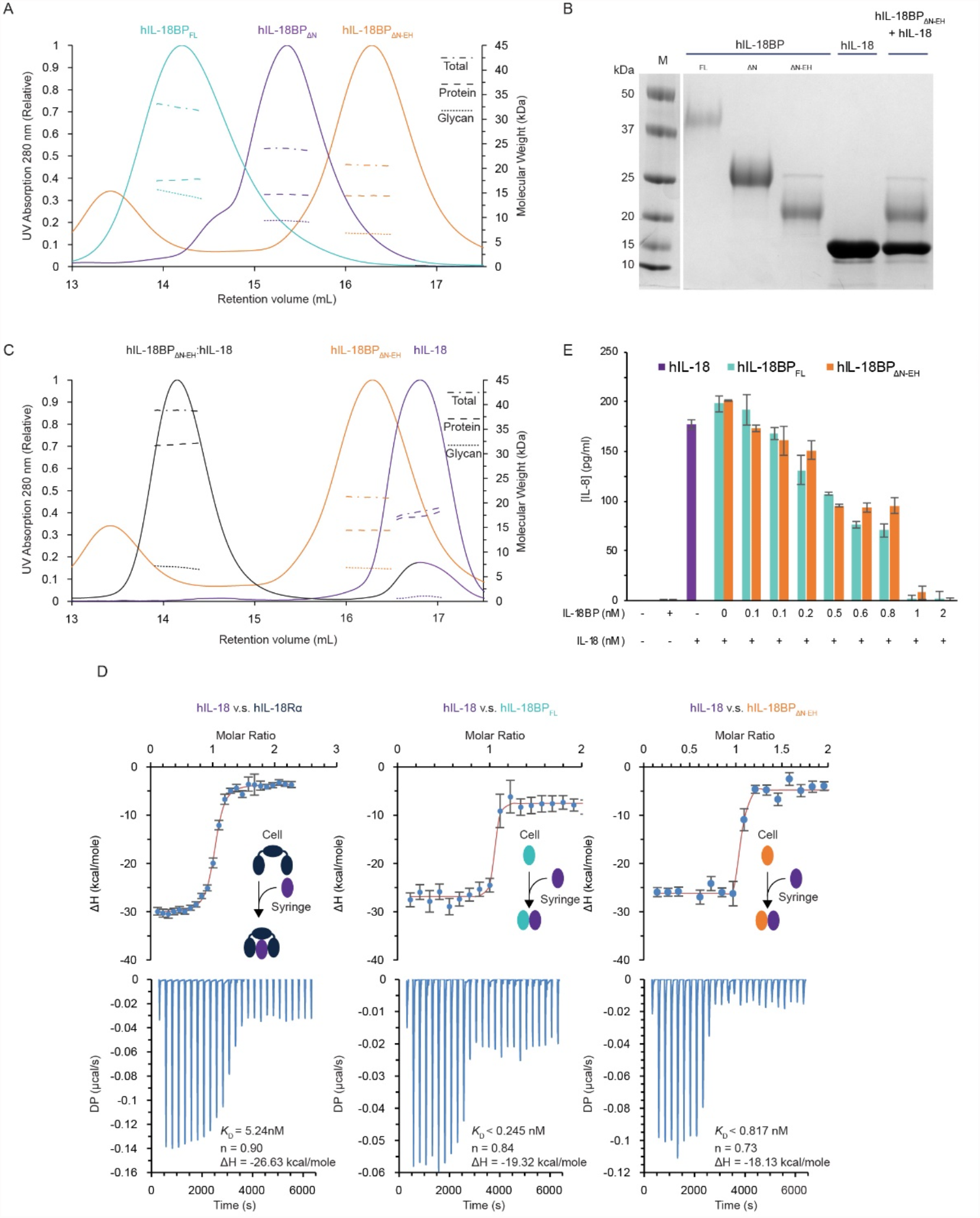
Biochemical characterization and functional activity of human IL-18 and IL-18BP. **(A)** SEC-MALLS analysis of hIL-18BP_FL_, hIL-18BP_ΔN_ and hIL-18BP_ΔN-EH_ protein (complex) respectively yielding less glycosylated and more homogenous material for crystallization purposes **(B)** SDS-PAGE of hIL-18BP_FL_, hIL-18_ΔN_ and hIL-18BP_ΔN-EH_ show decline in total protein and glycan mass caused by removal of O-glycosylated N-terminus and shaving of N-glycans by Endoglycosidase H (EH). SDS-PAGE of hIL-18 and hIL-18:hIL-18BP_ΔN-EH_ complex show presence and purity of both proteins. **(C)** SEC-MALLS analysis of hIL-18:hIL-18BP_ΔN-EH_ complex prior to crystallization allow accurate stoichiometry determination (A) & (C) Full lines show the SEC retention profile of samples (∼0.5-1 mg/ml) detected by UV (left axis, 280 nm). Interrupted lines show the molecular weights in kDa of the sample over the course of the SEC peak, split up by protein conjugate analysis in total (dashed and dotted), protein (dashed) and glycan (dotted) mass (right axis). **(D)** Bar plot of IL-8 levels in conditioned media after hIL-18 stimulation in presence or absence of hIL-18BP_FL_ or IL-18BP_ΔN-EH_. Recombinant IL-18 was pre-incubated for 1 h at 37°C with the indicated concentrations of full-length IL-18BP or IL-18BP_ΔN-EH_. KG-1 cells were left untreated (UT) or stimulated with IL-18 (purple) (0.5 nM final concentration) +/- IL-18BP (teal and orange), as indicated. After 18 h, IL-8 concentration in cell culture supernatants was measured by ELISA. Error bars display standard deviation (SEM). **(E)** Isothermal titration calorimetry (ITC) of hIL-18 in hIL-18Rα, hIL-18BP_FL_ and hIL-18BP_ΔN-EH_ shows recombinantly produced hIL-18 and hIL-18BP_ΔN-EH_ are both functionally competent. Bottom plots show the raw baseline-subtracted thermograms while top plots show integrated heats (blue dots) with fitted isotherm (red line). Diagrams in the isotherm plot show which protein was in the cell or syringe and what the expected final complex looks like.

To this end we produced hIL-18BP lacking the N- and/or C-terminal regions in HEK293S MGAT1 ^-/-^ cells. We found that only hIL-18BP truncated at the N-terminus (residues 63-194), hereafter termed hIL-18BP_ΔN_, could be sufficiently produced for structural studies. SEC-MALLS analysis of hIL-18BP_ΔN_ revealed the loss of 3 kDa of protein mass and 5.5 kDa of glycans (**Figure 1A** and **Supplemental Table** 1), confirming that the N-terminus of hIL-18BP is heavily glycosylated. Additional trimming of N-linked Man_5_GlcNAc_2_ glycans using Endoglycosidase H (Endo H) resulted in a further loss of 2.5 kDa of N-linked glycans (**Figure 1A, 1B**).

We initially verified the functional activity of hIL-18BP_ΔN_ treated with Endo H (IL-18BP_ΔN-EH_) in terms of its ability to sequester hIL-18 in stoichiometric fashion and high affinity. Indeed, SEC-MALLS analysis showed that hIL-18BP_ΔN-EH_ can establish a monodisperse complex with hIL-18 according to a 1:1 stoichiometry (**Figure 1C** and **Supplemental Table** 1). Using isothermal titration calorimetry (ITC), we compared the thermodynamic binding profiles and affinities of hIL-18BP_FL_, hIL-18BP_ΔN-EH,_ and the hIL-18 cognate receptor (IL-18Rα) to hIL-18 (**Figure 1D**). Our data show that hIL-18BP_ΔN-EH_ and hIL-18BP_FL_ are virtually indistinguishable from each other in their thermodynamic binding profiles and sub-nM affinities towards hIL-18. In contrast and consistent with previously published data (26), human hIL-18Rα binds to hIL-18 with lower affinity (K_D_=4.7 nM) than hIL-18BP. However, the binary complex affinity measured here is markedly higher than previously reported affinities (K_D_∼60 nM) (26) by surface plasmon resonance.

Finally, we interrogated the biological activity of hIL-18BP_ΔN-EH_ by using the hIL-18-responsive macrophage cell line, KG-1, the activity of which is manifested by NFκB signaling and pro-inflammatory cytokine and chemokine production (54). In this context, we measured the amount IL-8 as secreted from KG-1 cells upon hIL-18 stimulation, and found that both hIL-18BP_FL_ and hIL-18BP_ΔN-EH_ displays comparable inhibitory activities. (**Figure** 1E).

### Structural mimicry underlies the sequestration of human IL-18 by IL-18BP

Structural insights into the sequestration of human IL-18 by its cognate decoy receptor IL-18BP had remained elusive despite the growing importance of IL-18BP in physiology and disease. Using the truncated and glycan-shaved hIL-18BP_ΔN-EH_ we purified crystallization-grade hIL-18: hIL-18BP_ΔN-EH_ complex and determined the crystal structure of the complex to 1.8 Å resolution by molecular replacement using crystal structure of ectromelia poxvirus IL-18 binding protein (ectvIL-18BP) (23) (pdb 3f62) as a search model (**Figure** 2A, **Table** 1). Crystallographic refinement and the quality of the ensuing electron density maps were enhanced after correcting the X-ray data for data anisotropy using the STARANISO server (55). The crystal asymmetric unit contains one copy of the 1:1 complex and reveals how hIL-18BP uses the side of its h-type immunoglobulin β-sandwich fold to bind the β-trefoil structure hIL-18 (**Figure** 2A). hIL-18BP harbors two internal disulfide bridges connecting strands A and B (Cys86-Cys150), and B and F (Cys64-Cys89). In addition, three predicted N-glycan sites are occupied by an N-Acetylglucosamine residue (GlcNAc) at residues Asn79, Asn103 and Asn147. Our model for hIL-18BP starts at the beginning of the hIL-18BP_ΔN_ construct, i.e. Gln63, however no density was observed past Ala170 in the crystal solvent channels, consistent with the predicted disorder of the C-terminal tail (**Figure** 2C, **Figure** S1B). Human IL-18 features its signature β-trefoil fold and closely resembles the structure of unbound hIL-18 (pdb 3wo2) (all-atom r.m.s.d.=1.8 Å). Structure-based sequence alignments of human IL-18BP and structural superpositions against the two structurally characterized viral IL-18BPs, ectromelia virus IL-18BP (ectvIL-18BP) and yaba-like disease virus disease IL-18BP (yldvIL-18BP), and domain 3 of hIL-18Rα (hIL-18Rα_D3_) establish the strong conservation of the adopted fold, albeit with major variations in the CD loop mediating hIL-18 binding and AB loop (**Figure** 2B, **Figure** 2C, **Figure** 2D). Importantly, these analyses reveal that structural mimicry and direct steric competition underlie the sequestration of human hIL-18 by hIL-18BP (**Figure** 2C).

**Table 1.** Crystallographic data collection and refinement statistics.

|  | Human IL-18:hIL-18BP<sub>ΔN EH</sub> | Human IL-18:hIL-18BP<sub>ΔN EH</sub><br>Anisotropy Corrected |
| --- | --- | --- |
| DATA COLLECTION |  |  |
| Wavelength (Å) | 0.9762 | 0.9762 |
| Resolution range (Å) | 59.39 - 1.80 (1.87 - 1.80) | 59.39 - 1.80 (1.85 - 1.80) |
| Space group | C 1 2 1 | C 1 2 1 |
| Unit cell (a, b, c (Å), α, β, γ (°)) | 109.82 44.52 60.28 90 99.86 90 | 109.82 44.52 60.28 90 99.86 90 |
| Total reflections | 183218 (17814) | 140440 (6871) |
| Unique reflections | 26507 (2546) | 20705 (1035) |
| Multiplicity | 6.9 (7.0) | 6.8 (6.6) |
| Completeness (%) | 98.27 (96.15) | 77.2 (49.6) |
| Mean I/σ(I) | 15.67 (1.49) | 18 (2.9) |
| Wilson B-factor (Å<sup>2</sup>) | 36.35 | n.a. |
| R-meas | 0.053 (1.327) | 0.048 (0.680) |
| R-pim | 0.020 (0.495) | 0.018 (0.258) |
| CC1/2 | 0.999 (0.896) | 0.999 (0.898) |
| CC* | 1.000 (0.972) | n.a. |
| REFINEMENT (59.39 – 1.8 Å) |  |  |
| Reflections used in refinement | 26368 (2525) | 20691 (1306) |
| Reflections used for R-free | 2640 (248) | 2062 (130) |
| R-work/R-free** | 0.207 (0.531) / 0.230 (0.586) | 0.188 (0.264)/ 0.217 (0.320) |
| CC(work)/ CC(free) | 0.968 (0.860) / 0.959 (0.812) | 0.874 (0.689) / 0.858 (0.640) |
| Number of non-hydrogen atoms | 2309 | 2309 |
| macromolecules | 2136 | 2136 |
| ligands | 42 | 42 |
| solvent | 131 | 131 |
| RMS bonds (Å) / angles (°) | 0.011 / 1.34 | 0.011 / 1.34 |
| Ramachandran favored (%) | 98.05 | 98.05 |
| Ramachandran allowed (%) | 1.95 | 1.95 |
| Ramachandran outliers (%) | 0 | 0 |
| Rotamer outliers (%) | 0 | 0 |
| Clashscore | 3.48 | 3.48 |
| Average B-factor (Å<sup>2</sup>) | 34.68 | 34.68 |
| macromolecules | 34.03 | 34.03 |
| ligands | 66.67 | 66.67 |
| solvent | 35.15 | 35.15 |

Values in parentheses correspond to the highest resolution shell (1.87 - 1.80).
\*\*10% of reflections in R-free set

**Figure 2.**
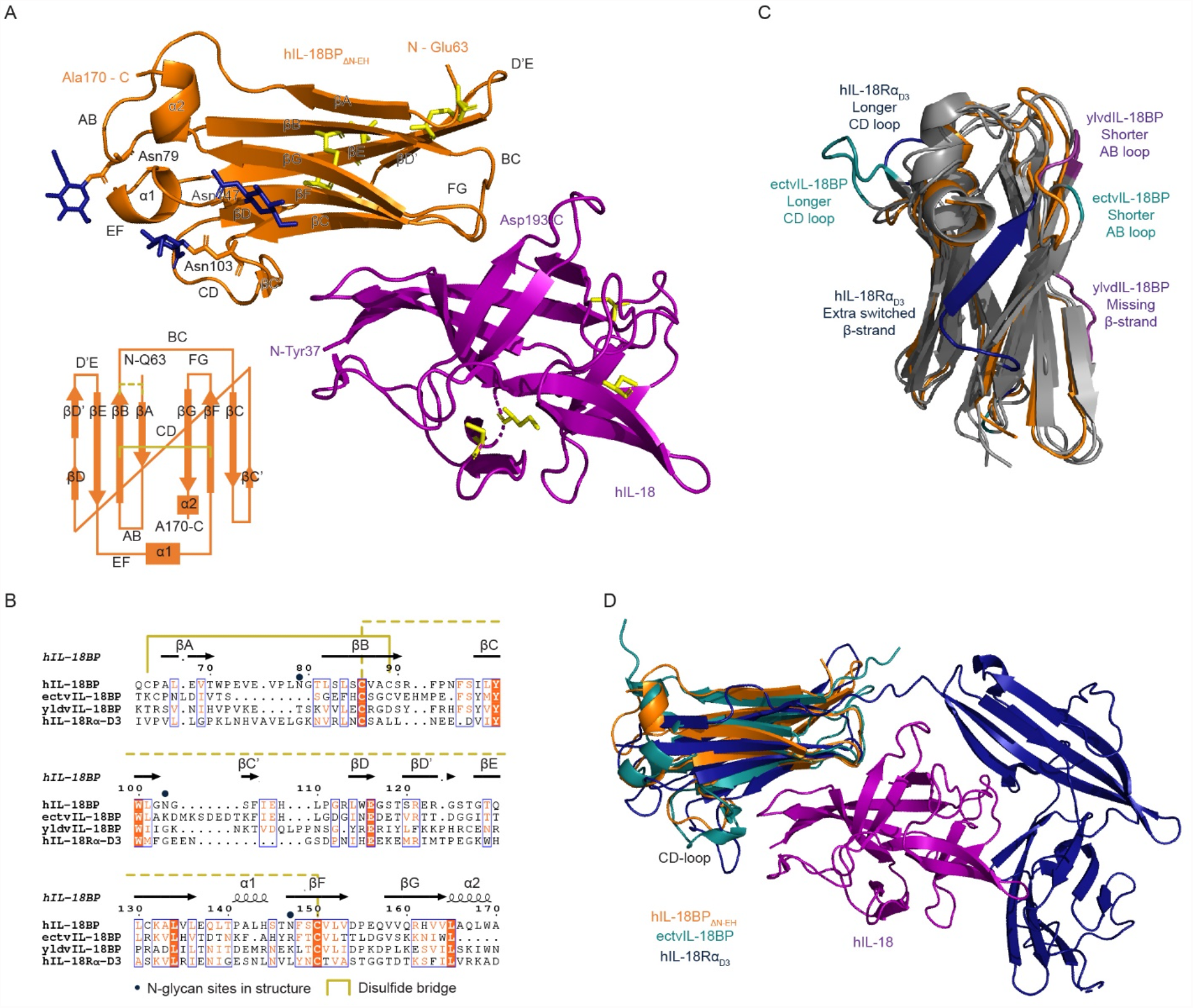
Crystal structure of human IL-18:IL18BP complex and comparisons with human IL-18 receptor and viral decoy receptors. **(A)** Cartoon representation of asymmetric unit of the crystal structure of hIL-18:hIL-18BP_ΔN-EH_ displaying hIL-18 (purple) interacting with hIL-18BP_ΔN-EH_ (orange). Asparagine residues occupied with GlcNAc (stick, blue) are shown in stick representation as well as all cysteines (yellow). Secondary structures of hIL-18BP_ΔN-EH_, β-strands, α-helices and loops, are labeled in order of occurrence. Additionally, a schematic of the Ig-fold of hIL-18BP_ΔN-EH_ is shown in the bottom left in which cysteine bridges are represented by yellow lines. **(B)** Structural sequence alignment based on existing crystal structures performed in Swiss-PdbViewer (72) and visualized by by ESPript (76) shows which residues spatially correspond to each other in the orthologous structures. Secondary structures are annotated by arrows (β-strand) and coils (α-helices) above the sequence. Linked disulfides are bridges by yellow lines (full or dashed). Asparagine residues occupied with GlcNAc in the model are labeled with blue dots. Residues with strict identity have an orange background, residues with similarity are colored orange. Groups of high similarity residues are surrounded by a blue box. **(C)** Crystal structure overlay of hIL-18BP_ΔN-EH_ (orange, pdb 7al7), ectvIL-18BP (teal, pdb 3f62(22)), yldvIL-18BP (purple, pdb 4eee(23)) and hIL-18Rα (dark blue, pdb 3WO4(26)) aligned to hIL-18BP_ΔN-EH_. Only notable differences on the level of secondary structures are colored accordingly, while stretches that align to hIL-18BP_ΔN-EH_ are left grey. Structural alignment of these structures shows that the core h-type Ig-fold is maintained with some strand and loop alterations, with an average all atom RMSD of 2.5 Å. Compared to human IL-18BP, ectv- and yldvIL-18BP have a shorter AB loop, while ectvIL-18BP has a significantly longer CD loop. **(D)** Cartoon representation of crystal structures of hIL-18BP_ΔN-EH_ (orange, pdb 7al7), ectvIL-18BP (teal, pdb 3f62(22)), yldvIL-18BP (purple, pdb 4eee(23)) and hIL-18Rα (dark blue, pdb 3WO4(26)) aligned to hIL-18 (purple, pdb 7al7).

Specifically, human IL-18BP covers the top of the hIL-18 β-barrel with one side of its β-sandwich scaffold composed of strands C, G and F and is slightly inclined to enable protrusion of loops FG and BC into the barrel. Additionally, residues from the CD loop and D strand mediate important interactions on the other side of the sheet (**Figure** 2A). In light of the plethora of structural information on human/viral IL-18BPs and human IL-18Rα in complex with hIL-18, curation and comparison of the observed interaction interfaces to distil key structure-function insights presents with considerable challenges. At the same time, annotation of amino acid positions that have been interrogated by mutagenesis would also be desirable. Together, such analyses could lead to a rationale for the exquisitely efficient sequestration of hIL-18 by hIL-18BP and might further fuel protein engineering approaches to modulate this sequestration potential. Therefore, we sought to develop a representation scheme that would allow effective parallel comparison of all structural data available together with mutagenesis data (**Figure** 3B). The binding interface can be subdivided into three sites: Site A consists of complementary hydrophobic patches on both hIL-18 and hIL-18BP. Sites B and C each engage a phenylalanine on hIL-18BP, Phe106 and Phe95 respectively, to protrude into a hydrophobic cavity on hIL-18 sealed by salt bridges and hydrogen-binding interactions (**Figure** 3A). Sequence alignments of hIL-18BP orthologs among vertebrates (**Figure** S2A) allows mapping of highly conserved residues on the surface of hIL-18BP that mediate sequestration of hIL-18, and shows that this region of hIL-18BP is the most conserved compared to any other part of the decoy receptor (**Figure** S2B). This suggests that the observed human IL-18BP:IL-18 complex can serve as a structural representative for all such complexes across species. When comparing the interface of human IL-18BP with its viral counterparts, these phenylalanine residues are largely conserved, with the exception of yldvIL-18BP, which contains a threonine instead of phenylalanine at site B. The most striking difference is the lack of salt bridges in viral IL-18BPs at site C. Conversely, human IL-18Rα does form these salt bridges but lacks the phenylalanine residues protruding into the hydrophobic cavity of hIL-18, while also lacking another hydrophobic substitute. At site B, hIL-18Rα also lacks the phenylalanine residue, but substitutes it for a methionine to fill the hydrophobic pocket (**Figure** 3B). Overall, site A is very similar for all interfaces, with mainly conservative substitutions of key interacting residues. The absence of these phenylalanine residues may explain the higher affinity of human IL-18BP to IL-18 than its receptor. Similar to IL-18Rα, but in contrast to the viral IL-18BPs, human IL-18BP exploits additional interactions via two side-chain-to-main-chain hydrogen bonds at site A.

**Figure 3.**
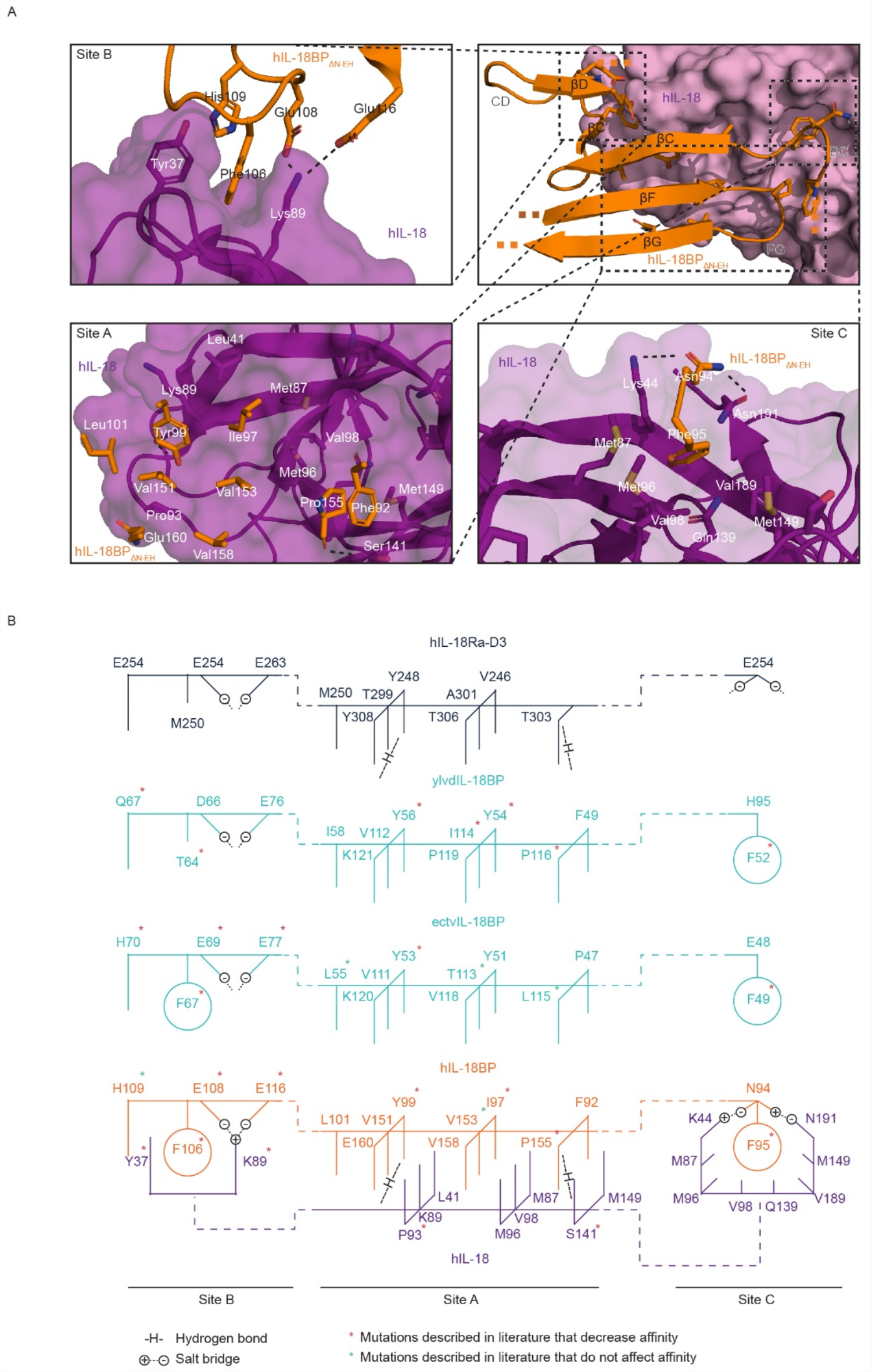
Interaction interface between human IL-18 and human IL-18BP and comparisons with human IL-18 receptor and viral receptor decoys. **(A)** Top right panel shows an overview of interacting structures of hIL-18BPΔN-EH (cartoon, orange) with hIL-18 (surface, purple). Structures of hIL-18BPΔN-EH shown consist out of the bottom half of the beta-sandwich, with the top half and non-interacting loops hidden. Top left, bottom left, and bottom right panels show zoomed-in views of the three main interaction sites B, A and C respectively, with hIL-18BPΔN-EH and hIL-18 shown in cartoon representation with a transparent surface overlay for hIL-18. Key residues are shown in stick representation colored by element (blue: nitrogen, red: oxygen, yellow: sulfur, white: polar hydrogens, orange/purple: carbon). Non-polar hydrogens are hidden. **(B)** Diagrams show simplified interactions at site A, B and C of hIL-18 (purple) with hIL-18BP (orange), viral ectvIL-18BP and ylvdIL-18BP (cyan), and domain 3 of hIL-18Ra (dark blue). The interaction between hIL-18 and hIL-18BP consists of 3 sites: A, B and C. Site A consists of interlocking rows of hydrophobic residues. Site B consist of a phenylalanine flanked by hydrophobic side chains supplemented with two salt bridges. Site C consists of a phenylalanine residing in a large pocket lined by hydrophobic residues supplemented by two salt bridges. Notable differences with ectvIL-18BP and yldvIL-18BP are missing salt bridges at site C, even though a capable side chain is present for ectvIL-18BP. ylvdIL-18BP also has a threonine replacing the phenylalanine at site B. hIL-18Ra-D3 does have the salt bridges at site C, but is lacking the phenylalanine with no substitution, while at site B, the phenylalanine is replaced with a methionine. Residues marked with a red asterisk have been reported to have reduced affinity after certain mutations. Residues marked with a green asterisk have been reported to not be affected by certain mutations. Mutagenesis data adopted from previously reported studies(22, 23, 26, 57, 84, 85).

### A disulfide-linked interaction site mediates a novel higher-order IL-18:IL-18BP complex

An intriguing feature of the IL-18–IL-18BP complex concerns an unexpected disulfide bridge between Cys74 of hIL-18 in the canonical IL-18:IL-18BP complex and Cys131 of IL-18BP participating in a symmetry-related complex. This results in a covalent tetrameric assembly with a 2:2 stoichiometry and twofold symmetry (**Figure** 4A). We note that such higher-order assembly was not observed in the preparatory stages of pre-crystallization samples as assessed by SEC-MALLS or SDS-PAGE (**Figure 1B** and 1C). However, the sample was concentrated to 30 mg/ml prior to crystallization. To verify whether such a higher order complex originated during this final protein concentration step, the concentrated sample was diluted back to 0.5 mg/ml and analyzed by SDS-PAGE and SEC-MALLS. SDS-PAGE shows a band at ∼38 kDa which can be resolved to the constituent protein components upon the addition of the reducing agent DTT prior to loading the sample, indicating the presence of a disulfide linked dimer (**Figure** 4B). We confirmed by mass spectrometry that both hIL-18 and hIL-18BP were present in this disulfide linked protein adduct, verifying a heterodimer as opposed to a homodimer of either. Additionally, SEC-MALLS shows that more than 90% of the previously concentrated sample now exists as a tetramer of 74 kDa, rather than the expected canonical IL-18:IL-18BP complex of 37 kDa (**Figure** 4C). Protein conjugate analysis of SEC-MALLS data validated the presence of both hIL-18BP and hIL-18 in the complex with a 2:2 stoichiometry, as any other combination of proteins and their glycans would not be compatible with their respective molecular masses.

**Figure 4.**
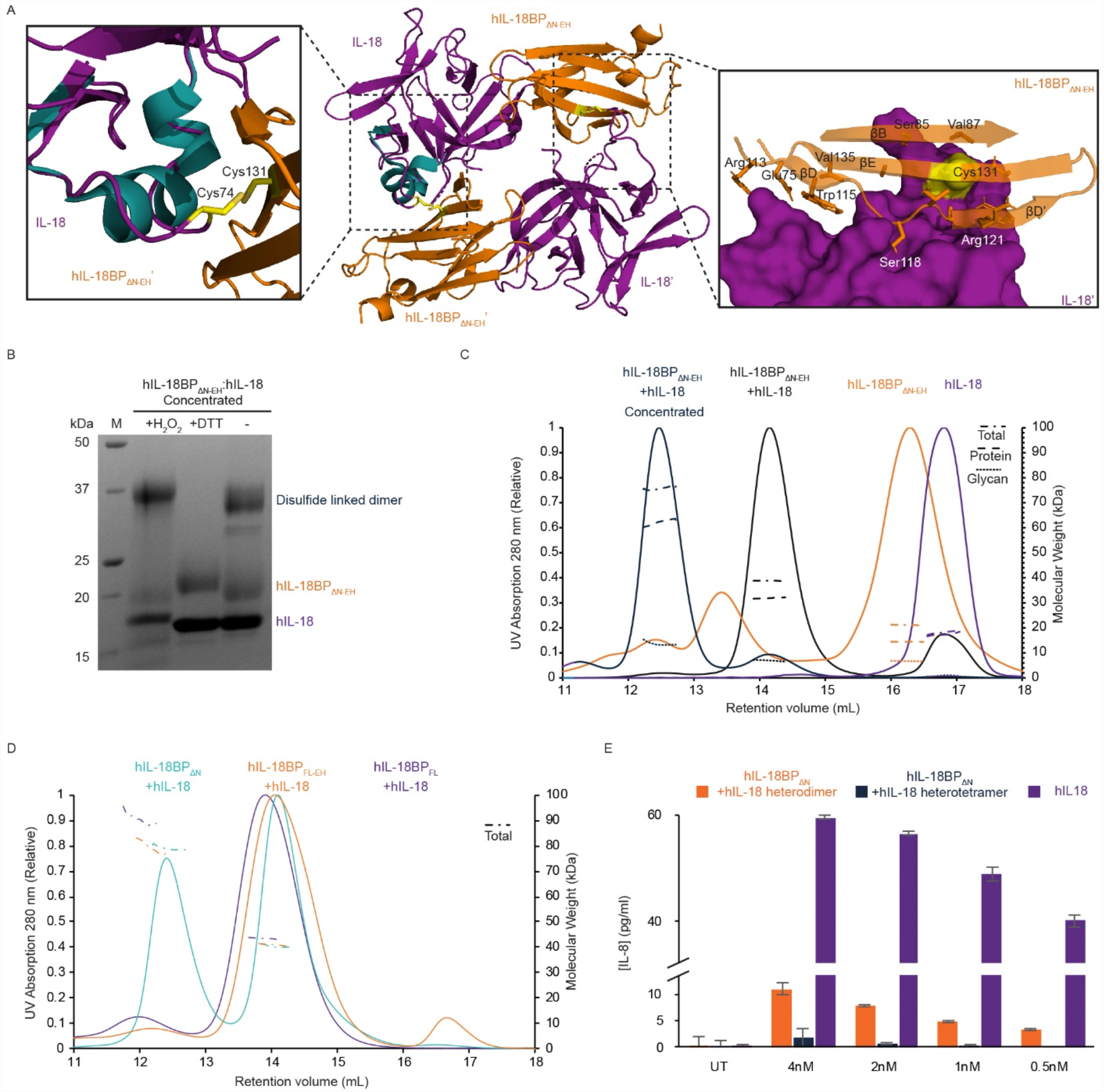
Tetrapartite assembly of human IL-18 and human IL-18BP. **(A)** Two complexes of hIL-18:hIL-18BPΔN-EH are linked through an intermolecular disulfide bridge between hIL-18 and hIL-18BPΔN-EH from neighboring symmetric related unit forming a heterotetrameric assembly with a novel interface. Middle panel shows an overview of the assembly using cartoon representation of hIL-18 (purple or teal) and hIL-18BPΔN-EH (orange) with the disulfide links involved shown in yellow stick representation. Left panel shows a zoomed-in view with an aligned overlay of the 310 α-helix from a crystal structure of unbound hIL-18 (teal, pdb 3wo2(26)) showing disruption of the helix upon formation of the disulfide link. The right panel displays the novel interface surrounding the disulfide link with hIL-18 shown in surface representation (purple, pdb 7al7) and hIL-18BPΔN-EH in cartoon representation (orange, pdb 7al7) with key residues displayed in stick representation. Strands, loops and key residues are labeled accordingly. **(B)** SDS-PAGE analysis of hIL-18:hIL-18BP_ΔN-EH_ complex sample after concentration to 30 mg/ml prior to crystallization and diluting back to 0.5 mg/ml with oxidizing (H_2_O_2_), reducing (DTT) or no agents added. Reduction results in dissociation of the otherwise disulfide-linked tetramer. **(C)** SEC-MALLS analysis of hIL-18, hIL-18BP_ΔN-EH_ and hIL-18:hIL-18BP_ΔN-EH_ complex before and after concentration to 30 mg/ml prior to and diluting back to 0.5 mg/ml. Line plots show the SEC retention profile of samples detected by UV (left axis). Interrupted lines plot the molecular weights of the sample over the course of the SEC peak, dissected by protein conjugate analysis in total (dashed and dotted), protein (dashed) and glycan (dotted) mass (right axis). **(D)** SEC-MALLS analysis of hIL-18:hIL-18BP_ΔN_ and hIL-18:hIL-18BP_FL_ complex sample after concentration to 30 mg/ml and diluting back to 0.5 mg/ml. Line plots show the SEC retention profile of samples detected by UV (left axis). Interrupted lines plot the molecular weights of the sample over the course of the SEC peak, dissected by protein conjugate analysis in total (dashed and dotted), protein (dashed) and glycan (dotted) mass (right axis). **(E)** Bar plot of IL-8 levels (y-axis, pg/mL) in conditioned media after human IL-18 stimulation in presence or absence of dimeric or tetrameric IL-18BP_ΔN-EH_:IL-18 complex. Recombinant IL-18 was pre-incubated for 1 h at 37°C with the indicated concentrations of full-length IL-18BP or IL-18BP_ΔN-EH_. KG-1 cells were left untreated (UT) or stimulated with IL-18 (purple) (0.5 nM final concentration) +/- IL-18BP (teal and orange), as indicated. After 18 h, IL-8 concentration in cell culture supernatants was measured by ELISA. Error bars display standard deviation (SEM).

Interestingly, in the structure of unbound hIL-18 and when in complex with its decoy and signaling receptors, the cysteine involved in the observed novel disulfide bond with IL-18BP is part of a 3_10_ helix and is not surface exposed (**Figure** 4A, left panel). This region in inbound IL-18 is rather amphipathic in terms of physicochemical properties and interacts with several detergent molecules (CHAPS) in pdb 3wo2 (26). Thus, to participate in the observed disulfide bond, the cysteine has flipped outwards and is accompanied by a structural rearrangement thereby disrupting the helical structure (**Figure** 4A, left panel). Interestingly, an ensemble of structural models of IL-18 determined by NMR has shown this helix to be very flexible (56), which would favor such dynamics. We note that for the reported crystals structure of hIL-18 complexed with ectvIL-18BP and yldvIL-18BP (pdb 3f62 and 4eee respectively), all free cysteines were mutated to serine, thereby excluding the possibility of such a disulfide linkage, although this particular helix/loop segment was not modelled due to its apparent flexibility. Surrounding the cysteine disulfide bridge is a novel interface mediated by the beta strands E, B, D and D’ of the opposite side of the beta-sandwich compared to the canonical interface and is maintained by a several hydrophobic residues on both molecules.

Because this hetero-tetramer was only initially observed with IL-18BP_ΔN-EH_ lacking the N-terminal and N-glycans, we also investigated if a similar higher-order complex could be formed with IL-18BP_FL_ or endogenously glycosylated IL-18BP_ΔN_. Full-length IL-18BP was complexed with IL-18, concentrated to 0.5 mg/mL and analyzed by SEC-MALLS. No tetrameric complex was observed using IL-18BP_FL_, either in its glycosylated or deglycosylated form, suggesting that removal of the N-terminal segment of IL-18BP promotes the formation of this hetero-tetrameric complex (**Figure** 4C). We note that Cys51 is part of the truncated and likely pairs up with Cys131 in the full-length protein, thus preventing heterotetramer formation. Endogenously glycosylated IL-18BP_ΔN_, without Endo H treatment, also formed a tetrameric complex with IL-18 at high concentrations, albeit at a lower rate than IL-18BP_ΔN-EH_ (**Figure** 4D).

The presence of a disulfide bridge linking IL-18BP_ΔN_ and IL-18 distal to the binding interface in the hetero-tetrameric assembly observed here may result in better sequestration and inhibition of IL-18 than in the canonical 1:1 complex. Furthermore, the site where disulfide-linked IL-18BP engages on IL-18 would also sterically clash with the eventual binding site of IL-18 to domain 1 of IL-18Rα. To test this, both the heterodimeric or hetero-tetrameric complex of IL-18BP_ΔN-EH_ and IL-18 was purified and used to stimulate KG-1 cells. While both the dimeric and tetrameric complex inhibited IL-18 activity compared to IL-18 alone, the tetrameric assembly restrained IL-18 activity more effectively than the dimeric complex, resulting in less inflammatory cytokine production from KG-1 cells (**Figure** 4E).

## Discussion

Over the past two decades and in the absence of an experimentally determined structure for the human IL-18–IL-18BP complex, numerous studies have used orthogonal methods to predict how human IL-18BP interacts with human IL-18 to efficiently sequester and prevent signaling. To date, structural features and binding interfaces between human IL-18 and human IL-18BP have been inferred from computational models (44), human and viral mutagenesis studies (45–49) and homologous structures of viral IL-18BPs in complex with human IL-18 (22, 23). Additionally, the ternary assembly of human IL-18 in complex with its cognate receptors, IL-18Rα and IL-18Rβ (26), has shed some light on the potential for competitive binding of human IL-18BP. The high-resolution structural analysis of human IL-18BP bound to IL-18 presented here is imperative to further understand and exploit the ultra-high affinity binding mode that characterizes this unique decoy receptor-like protein (22, 23). Human IL-18BP binds the same epitope of hIL-18 used by viral IL-18BPs to compete with the third ectodomain of hIL-18Rα. The majority of the interface (Site A) consists of two large, complementary hydrophobic patches, mimicking the landscape of hydrophobic peaks and troughs on each side of the interface, supplemented with two side-chain-to-main-chain hydrogen bonds. Comparing the structure of hIL-18BP with its viral orthologs confirms that this large hydrophobic patch on site A is highly conserved. yldvIL-18BP contains some additional hydrophobic residues (Ile78 and Leu80) on the DE loop interacting with hIL-18 (Leu 5 and Glu6) at the perimeter of the interface. In all other structures, except yldvIL-18BP, this loop is constrained as an additional βD’ strand. On each side of the hydrophobic patch, a phenylalanine on hIL-18BP is tightly inserted inside hydrophobic cavities on hIL-18 and additionally locked down by salt bridges (Site B and C, **Figure** 3B). Mutating either phenylalanine residues to alanine reduces the ultra-high picomolar affinity of hIL-18BP down to low-nanomolar affinity (22, 23, 57). Mutations of the corresponding residues in viral IL-18BPs have a similar effect, highlighting the importance of these specific residues for binding. Interestingly, in hIL-18Rα the corresponding residue is either substituted by methionine (Site B) or completely lacking (Site C), although site B is flanked by additional hydrophobic residues residing on hIL18Rα’s longer CD’ loop. The absence of these seemingly critical phenylalanine residues may, in part, explain the higher affinity of hIL-18BP to IL-18 than its receptor. Similarly, mutating Glu108 and Glu106 on hIL-18BP involved in the salt bridges at Site B accordingly resulted in a lower affinity, as well as mutating Lys53 on hIL-18 which is located on the other end of the salt bridges (58, 59). Mutations in Site C particularly have not been reported, because in both viral structures the corresponding residues (Glu48 in ectvIL-18BP and His51 in yldvIL-18BP) are not involved in salt bridge formation. Overall, it is clear that hIL-18BP utilizes a unique constellation of key residues, contributing to ultra-high affinity binding to IL-18, however further investigation is required to determine binding hotspots important for this interaction. Whether or not the lower affinity of viral IL-18BPs is actually advantageous or disadvantageous for their function is unclear.

Unexpectedly, we observed a higher order assembly of IL-18 and IL-18BP in our crystallographic model, coordinated by an otherwise buried cysteine in IL-18, disrupting an alpha helix in the process. Using SEC-MALLS and mass spectrometry, we confirmed the formation of a hetero-tetramer, consisting of two complexes of IL-18:IL-18BP, with an additional interaction interface distal to the canonical binding site of IL-18BP. Interestingly, this hetero-tetrameric complex restrained IL-18 activity more effectively than the canonical dimeric 1:1 complex. Given that high concentrations and removal of the N-terminal region appear to be required, it is likely not biologically relevant. However, this fortuitous observation suggests that this interface may be amenable to further engineering to increase the sequestration potency of IL-18BP, possibly in combination with domains 1 and 2 of IL-18Rα. It brings to light plasticity in IL-18, the possible reactivity of Cys131 in IL-18BP, and the possible utilization of a hydrophobic patch at opposite ends to the IL-18:IL-18BP interface to create a new interaction interface. An analogous yet distinct bivalence has been observed in viral yldvIL-18BP. This ortholog lacks a highly conserved phenylalanine residue in site B, however it compensates for this by forming a 2:2 complex with IL-18, driven by disulfide-linked IL-18BP homodimerization (23). However, it is important to note that the 2:2 hetero-tetrameric complex seen here for human IL-18BP engages a separate interface, and is driven by an additional interaction with IL-18 cytokine.

The structural model presented here will facilitate further the structure-driven development of novel therapeutics to either disrupt or mimic hIL-18BP binding. Aberrant IL-18 signaling has been implicated in numerous inflammatory diseases including RA, AOSD, SLE, sickle cell disease, and more recently, the inflammatory phase of coronavirus infection. (28, 29, 36, 37, 60). Coronavirus disease (COVID-19), caused by the SARS-CoV2 virus, displays a broad range of clinical symptoms, resulting in disease severity extending from mild to fatal outcomes. Accumulating evidence has shown that tissue damage in the later stages of severe COVID-19 infection is driven by cytokine release syndrome (CRS), also known as cytokine storm (38, 39, 61). Recent studies have detected elevated levels of IL-18 in patients with moderate and severe COVID-19 infection, which strongly correlated with disease severity (38, 39, 62). Longitudinal analysis of patients infected with SARS-CoV-2 showed IL-18 levels remained elevated in severe COVID-19 patients admitted to ICU, while cytokine levels declined over time in moderately affected patients. Moreover, both IL-18 and IL-18BP have been identified as promising biomarkers to predict disease severity and likelihood of death after SARS-CoV-2 infection (38, 63). Elevated levels of IL-18 were also detected in multisystem inflammatory syndrome (MIS) and Kawasaki syndrome in children following SARS-CoV-2 infection, and may be used as a biomarker to distinguish these diseases from similar hyperinflammatory syndromes (37). Increased IL-18 production has also been associated with activation of cytotoxic mucosal-associated invariant T (MAIT) cells during coronavirus infection, leading to lung tissue damage and increased disease severity (40). These studies suggest that modulation of IL-18 activity may represent a novel therapeutic opportunity for COVID-19, however, no clinical trials with recombinant IL-18BP or IL-18-specific antagonistic have yet been initiated (64, 65). However, because of the pleiotropic activity of IL-18, there will likely be a narrow therapeutic window to exploit the beneficial activity of IL-18 for effective viral clearance, while avoiding aberrant inflammation and tissue damage associated with high IL-18 activity during late-stage infection. In addition, IL-18 has been shown to display potent anti-tumor activity, however clinical trials using recombinant IL-18 have shown limited efficacy (42). IL-18BP, produced in the tumor microenvironment, has been identified as an ‘immune checkpoint’, hampering the therapeutic application of recombinant IL-18 in cancer (66). Directed evolution of yeast-displayed mouse IL-18 variants identified a decoy-resistant IL-18 that retains signaling activity but escapes inhibition by IL-18BP. This IL-18 variant displayed potent cytotoxic T cell activation, NK cell maturation and tumor growth inhibition compared to its wildtype counterpart (66). The high-resolution structural models presented here, in particular the experimentally determined interaction interface of IL-18 and human IL-18BP, will be of crucial importance for additional protein engineering and future drug design for the development of novel therapeutics for use in viral infections, autoimmune and inflammatory diseases and cancer.

## Materials and Methods

### Plasmids, protein expression constructs and cell Lines

All constructs were created by a traditional restriction ligation approach. Restriction enzymes, T4 ligase and Q5 polymerase were purchased from NEB (New England Biolabs, Ipswich, Massachusetts, USA). All primers were purchased from IDT (Integrated DNA Technologies, Leuven, Belgium). Recombinant DNA was purchased from GenScript (GenScript, Piscataway, New Jersey, USA). The mature sequence of human IL-18 (residues 37-193, UniProt ID: Q14116) was codon optimized for expression in *E. coli* and purchased in the pUC57 vector from GenScript (GenScript, Piscataway, New Jersey, USA). The sequence was cloned into the pET42a plasmid (Cat No 70561, Novagen, Merck, Overijse, Belgium) in frame with an N-terminal caspase3-cleavable glutathione S-transferase (GST)-tag (26). The introduction of an extra N-terminal hexahistidine (His_6_)-tag was performed by polymerase chain reaction (PCR). The sequence of human IL-18BP (residues 1-194, UniProt ID: O95998) was purchased in the pUC57 vector at GenScript (GenScript, Piscataway, New Jersey, USA). The sequence was cloned in frame with a C-terminal caspase3 site followed by an AviTag and a His_6_ tag. The construct was further cloned in frame with an N-terminal chicken RTPµ-like signal peptide sequence (67) that replaced the native signal peptide sequence (residues 1-30). For crystallization purposes, the N-terminus (residues 31-60) was removed by PCR (IL-18BP_ΔNΔN_) and the construct was cloned in the same pHLSec plasmid containing an N-terminal chicken RTPµ-like signal peptide sequence and a caspase 3 cleavable Avi-His_6_-tag. All constructs were validated by Sanger sequencing by GATC BioTech (Konstanz, Germany) before further experiments were performed.

### Expression and purification of recombinant Proteins

BL21(DE3) *E. coli* transformed with plasmid pET42 expressing N-terminally GST- and HIS-tagged human IL-18 were grown at 37°C in Lysogeny broth (LB) medium containing kanamycin (25 μg/ml) as selection marker for the pET42a plasmid (Cat No 70561, Novagen, Merck, Overijse, Belgium). When the optical density at 600 nm (OD600) reached 0.6, the expression of human IL-18 was induced by addition of isopropyl-β-D-thiogalactopyranoside (ITPG) at a final concentration of 1 mM, after which the culture was incubated at 28°C for 5 hours. The bacteria were harvested by centrifugation (7,000 × g for 10 min at 4°C) and the cellular pellet was stored at −80°C. The bacterial pellet was thawed and resuspended in HEPES buffered saline (HBS, 20 mM HEPES, 150 mM NaCl, pH 7.4) with DNaseI (REF 10104159001, Roche Diagnostics, Vilvoorde, Belgium). The cells were lysed by sonication with a Qsonica macrotip sonicator (Newtown, Connecticut, USA) (on-time 4 min; pulse on 30 sec; pulse off 30 sec; amplitude 70%) while cooled on ice. The lysate was clarified by centrifugation (20,000 × g for 45 min at 4°C) and filtration using a 0.22-μm Millipore™ Steritop™ Sterile Vacuum Bottle-Top Filter (Thermo Fisher Scientific, Merelbeke, Belgium), and loaded onto a HisTrap^™^ HP 5 ml column (GE Healthcare, Diegem, Belgium) equilibrated with HBS. The column was washed (HBS, 50 mM imidazole) before eluting the protein (HBS, 250 mM imidazole) following desalting using a HiPrep™ 26/10 Desalting column (GE Healthcare, Diegem, Belgium) to remove imidazole. The His_6_-GST-tag was removed by caspase 3 (produced in-house) cleavage at 37°C for 1 hour. To remove the uncleaved protein and the His_6_-GST-tag, the sample was again loaded onto a HisTrap^™^ HP 5 ml column (GE Healthcare, Diegem, Belgium). The flow-through was collected and concentrated before injection onto a HiLoad® 16/600 Superdex® 75 pg column (GE Healthcare Diegem, Belgium) using HBS as running buffer. Fractions containing human IL-18 were pooled, flash frozen and stored at -80°C. The purity of the protein was evaluated on SDS-PAGE stained with Coomassie blue.

### Expression and purification of recombinant human IL-18BP

Adherent HEK293S MGAT1-/- cells (68) were grown in 5-layer cell culture flasks (Falcon® Multi-Flask, Corning, New York, USA) in Dulbecco’s Modified Eagle Medium (DMEM) (Gibco^™^, Life Technologies, Thermo Fisher Scientific, Merelbeke, Belgium) supplemented with 10% Fetal Calf Serum (FCS) (Bodinco, Alkmaar, Netherlands). Upon transfection, the growth medium was exchanged for DMEM supplemented with 3.6 mM valproic acid (item 13033, Cayman Chemical Company Europe, Tallinn, Estonia). Transient expression of human IL-18BP or IL-18BP_ΔNΔN_ was achieved using branched polyethyleneimine (PEI) (Mn ∼ 10 000, Cat.:40,872-7, Sigma-Aldrich, Diegem, Belgium) as transfection reagent (67). After 4 days of expression, filtered conditioned medium was loaded onto a 5 ml cOmplete^™^ His-tag Purification Column (Roche Diagnostics, Vilvoorde, Belgium). The protein was eluted with 250 mM imidazole in HBS after which the imidazole was removed using a HiPrep™ 26/10 Desalting column (GE Healthcare, Diegem, Belgium). The Avi-His_6_-tag was cleaved by caspase-3 (produced in-house) and the glycans were trimmed by EndoH (produced in-house) at 37°C for 1 hour. The flow-through of the HisTrap^™^ HP 5 ml column (GE Healthcare, Diegem, Belgium) containing the cleaved protein was collected, concentrated and injected onto a HiLoad® 16/600 Superdex® 200 pg column (GE Healthcare Life Sciences) using HBS as running buffer. Fractions containing human IL-18BP or IL-18BP_ΔNΔN_ were pooled, flash frozen and stored at -80°C. The purity of the protein was evaluated on SDS-PAGE stained with Coomassie.

### Multi-Angle Laser Light Scattering

Purified protein samples at a concentration of 0.5 mg ml^-1^ were injected onto a Superdex Increase 10/300 GL column (GE Healthcare, Diegem, Belgium) that was pre-equilibrated with HBS. The column was coupled in line with a UV-detector (Shimadzu, Brussels, Belgium), a miniDAWN TREOS multi-angle laser light scattering (MALLS) detector (Wyatt, Santa Barbara, California, USA) and an Optilab T-rEX refractometer (Wyatt, Santa Barbara, California, USA). Refractive index (RI) increment values (dn/dc) of 0.185 ml g^-1^ and 0.155 ml g^-1^ were used for protein and glycan analysis, respectively. Bovine serum albumin (BSA) (Pierce, Merelbeke, Belgium) was used as standard to correct for band broadening. The resulting data was analyzed using the ASTRA6.1 software (Wyatt, v6.1) and errors were calculated in Microsoft Excel.

### Isothermal Titration Calorimetry

All proteins were buffer matched in HBS from the same batch using size exclusion chromatography and all the solutions were degassed. Protein concentrations were determined with the NanoDrop^™^ 2000 (Thermo Fisher Scientific, Merelbeke, Belgium) using their corresponding extinction coefficients (absorbance of 1%). Experiments were carried out on the MicroCal VP-ITC calorimeter (Malvern Panalytical, Malvern, United Kingdom) at 37°C. Titrations were always preceded by an initial injection of 3 μl, and were carried out using 10 μl injections applied 300 s apart. The sample was stirred at a speed of 300 r.p.m. throughout. The data was recorded using the Origin® scientific plotting software (Version 7.0, MicroCal, Malvern Panalytical, Malvern, United Kingdom) and further analyzed using NITPIC (69) and SEDPHAT (70) fitting the data with a “one to one” binding model.

### Bioactivity assays

KG-1 cells were cultured in RPMI media (Gibco) supplemented with 10 % fetal calf serum (Bodinco). Cells were cultured at 37°C in a humidified atmosphere with 5 % CO_2_. KG-1 cells were plated at 2 × 10^5^ cells per well in 24-well plates. Recombinant IL-18 was pre-incubated with IL-18BP_FL_,IL-18BP_ΔN-EH_ or complexes thereof as indicated, for 1 h at 37 °C. KG-1 cells were left untreated or stimulated with IL-18 (0.5 nM final concentration) +/- IL-18BP, as indicated. Cells were incubated with stimuli for 18 h and cell culture supernatants were subsequently collected. Culture supernatants were centrifuged for 5 minutes at 1000 g to remove cells and frozen at -80°C.

### Detection of cytokines by ELISA

Human IL-8 was measured from cell culture supernatants using commercially available specific paired ELISA kits (R&D Systems). Briefly, a 96 well plate was coated with 50 ml of capture antibody per well according to manufacturer’s instructions, and incubated overnight at room temperature. Wells were washed three times with 150 ml wash buffer (0.05% Tween-20 in PBS, pH 7.2), then blocked for 1 h in reagent diluent (1% BSA in PBS, pH 7.2). Samples were diluted as required in reagent diluent and 50 ml of each sample was added to wells and incubated for 2 h at room temperature. Appropriate standards were prepared for each ELISA assay according to manufacturer’s instructions. Wells were washed again three times, followed by 50 ml of biotin-conjugated detection antibody incubated for 2 h. Wells were washed three times and 50 ml streptavidin-HRP was added to wells and incubated for 30 minutes in the dark. Finally, wells were washed three times, followed by addition of 50 ml TMB substrate solution. The reaction was stopped with 30 ml of 2M sulfuric acid and absorbance was read at 450 nm on an ELISA plate reader (Fluostar). All cytokine assays were carried out using triplicate samples from each culture.

### Bioinformatic analyses

All sequence alignments were performed by Clustal Omega (71). Structural alignments were performed in Swiss-PdbViewer (72). Glycan prediction were performed using NetNGlyc (73) and NetOGlyc servers (74). Protein disorder was predicted using IUPred (75). Sequence alignments were visualized by ESPript (76). Protein structures were visualized in PyMOL (77, 78). Interaction interface was inferred using the PISA server (79).

### Crystal structure determination and refinement

During the purification of IL-18BP_ΔN_, purified IL-18 was added in a 1:2 ratio prior to caspase 3 and Endo H digests. The complex was purified via SEC and concentrated to 35 mgml^-1^. Sitting-drop vapor diffusion experiments were set up in Swissci 96-well triple drop plates (Molecular dimensions, Suffolk, United Kingdom) with commercial sparse matrix crystallization screens. Using a Mosquito® liquid handling robot (TTP Labtech, Hertfordshire, United Kingdom), the protein was mixed with mother liquor in a 1:1 ratio in a final volume of 200 nl. The plates were incubated at 20°C. An initial hit in the Crystal Screen (Cat No HR2-110, Hampton Research, Aliso Viejo, California, USA) in condition D10 (0.2 M Calcium acetate hydrate, 0.1 M Sodium cacodylate trihydrate pH 6.5, 18% w/v Polyethylene glycol 8000) was optimized by varying pH (6.2-7.1) and PEG concentration (12%-27%). Crystals obtained from the optimization screen were used for seeding new plates, finally yielding single crystals. These crystals were cryoprotected in mother liquor supplemented with 20% glucose prior to cryo-cooling in liquid nitrogen. Diffraction data was initially collected under cryogenic conditions at a synchrotron radiation source, namely beamline PETRA III in Hamburg.

The diffraction data were integrated using XDS (80) and were treated for anisotropy using the STARANISO server (55). Initial phases were determined by maximum-likelihood molecular replacement in PHASER (81) using a search model derived from the structure of hIL-18 in complex with ectvhIL-18BP (pdb 3f62). Iterative cycles of structure building and refinement were performed in COOT (82) and PHENIX.refine (83) respectively.

Structure coordinates and structure factors for the human IL-18:IL18BP complex have been deposited in the Protein Data Bank (PDB) under accession code 7al7.

## Acknowledgements

We thank AB2 Bio Ltd. (Lausanne, Switzerland) for clinical grade recombinant IL-18BP and Prof. Seamus Martin, Trinity College (Dublin, Ireland), for providing KG-1 cells. J.A., Y.B. and D.C. are research fellows of Research Foundation Flanders (FWO), Belgium. This project has received funding from the European Union’s Horizon 2020 research and innovation program under grant agreement No 779295.

## Author Contributions

S.D. and J.A. carried out recombinant protein production and purification, biophysical studies, and structural studies with contributions from S.N.S. D.C. carried out bioactivity assays. Y.B. provided input on the generation of the glyco-engineered hIL-18BP variant. A.S., E.J.S., and C.G. provided initial research materials. S.D., J.A., D.C. and S.N.S. wrote the manuscript with contributions from all authors. S.N.S. conceived and supervised the project.

## Supplementary Information for

**Supplemental Figure 1.**
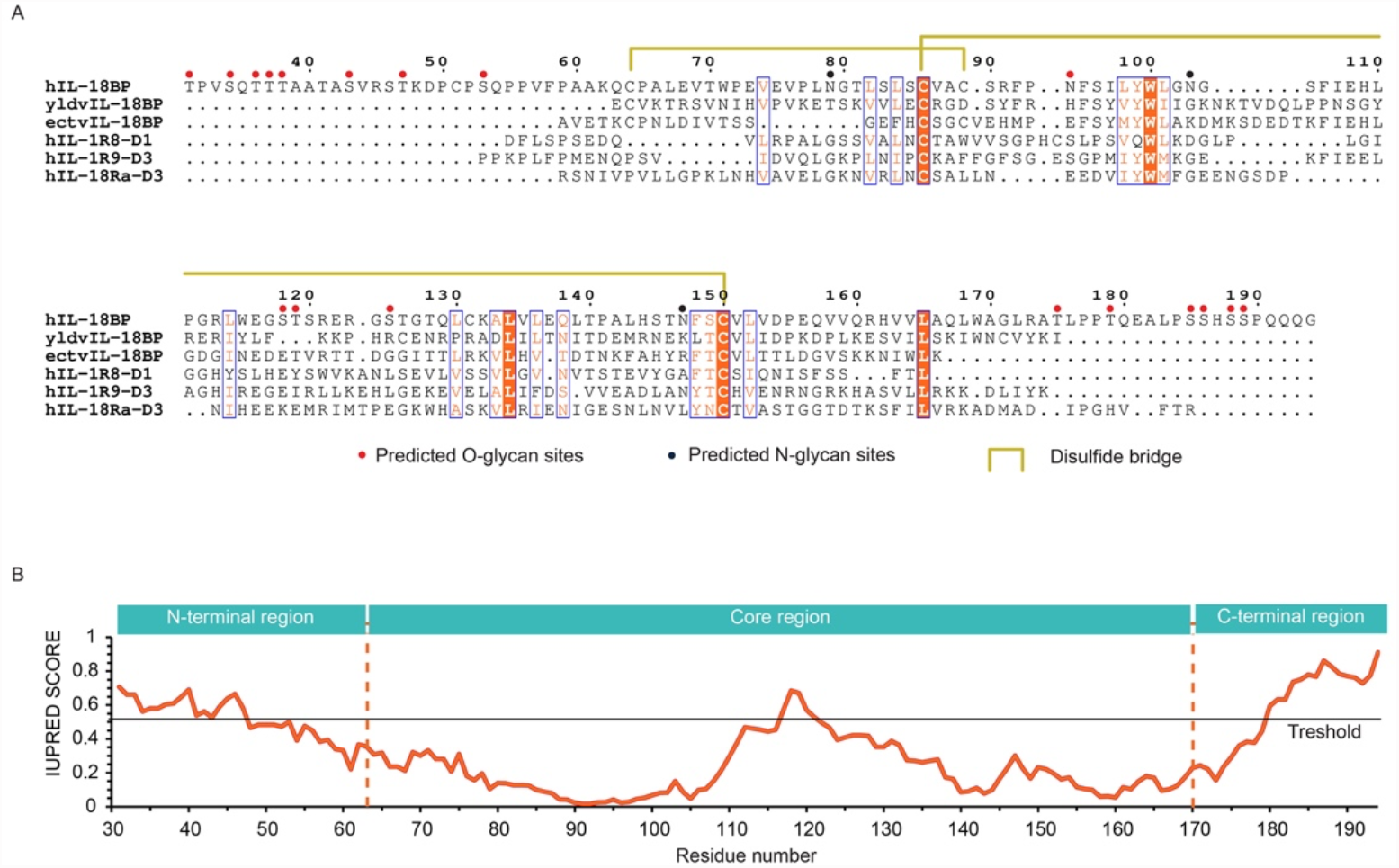
Sequence analysis for protein construct optimization. **(A)** Sequence alignment performed by Clustal Omega^1^ visualized by ESPript^2^ of hIL-18BP and related sequences. The core-region was delineated based on the sequence alignment, disordered region prediction and existing crystal structures of viral IL-18BPs. Gaps in the sequence alignment are represented by dots. Residues with strict identity have an orange background, residues with similarity are colored orange. Groups of high similarity residues are surrounded by a blue box. Predicted N-^3^ and O-glycan^4^ sites are displayed as blue and red dots respectively. Linked disulfides are represented by yellow lines (full or dashed). **(B)** IUPRED prediction^5^ of disordered regions in human IL-18BP show that the extended N- and C-termini are likely disordered and flexible. Disorder threshold is represented by a black horizontal line. Delineation of tested terminal truncations is represented by vertical dashed orange lines.

**Supplemental Figure 2.**
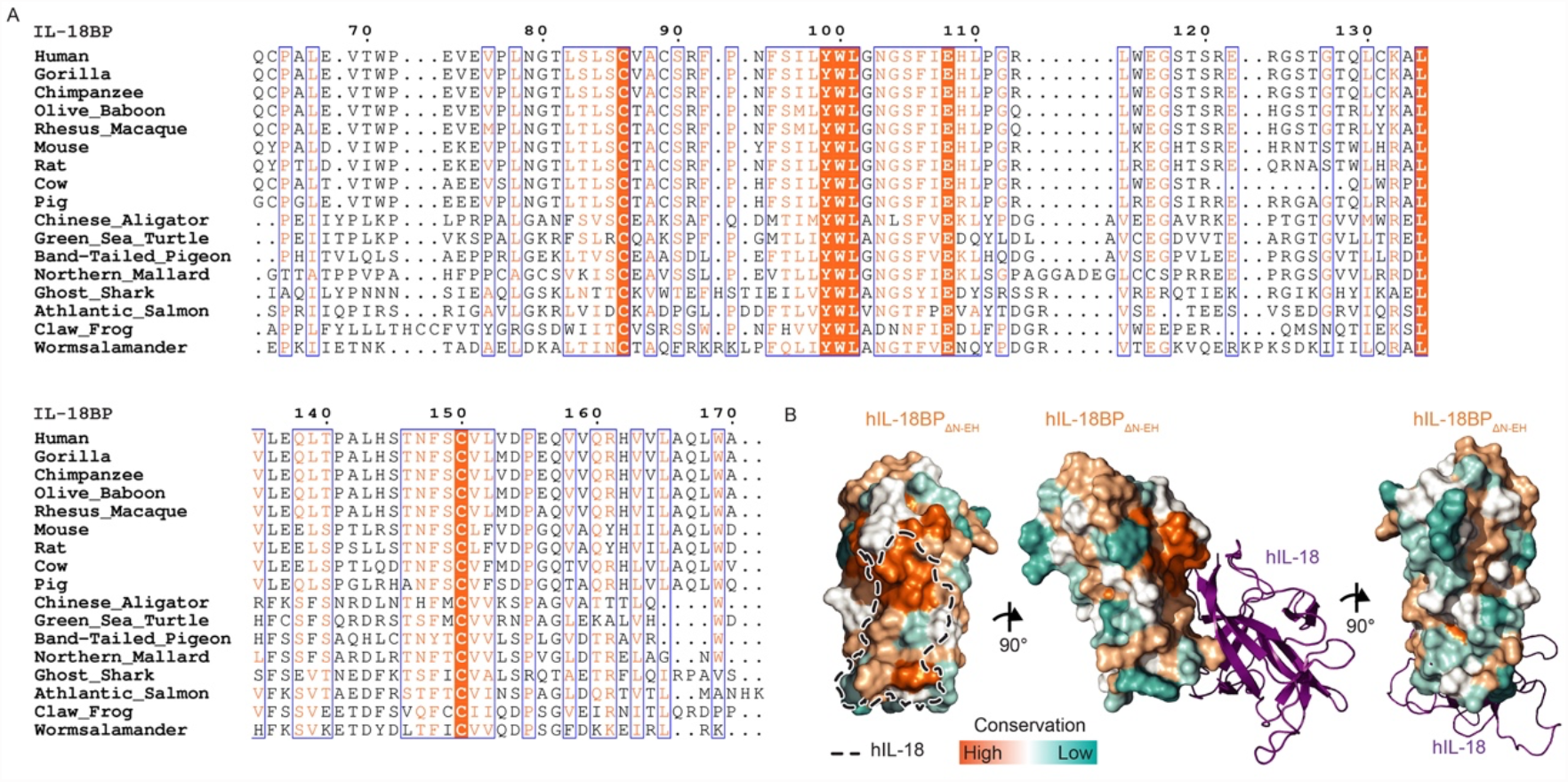
Sequence conservation for hIL-18BP. **(A)** Sequence alignment of core region of orthologous IL-18BPs between species across the animal kingdom. Residues with strict identity have an orange background, residues with similarity are colored orange. Groups of high similarity residues are surrounded by a blue box. Color scale above the sequence of hIL-18BP represents conservation levels determined by ConSurf^6^, with high conservation colored in orange, low conservation levels in teal and white in between. The same scale is used in panel B. **(B)** Surface representation of hIL-18BP_ΔN-EH_ colored by residue conservation. hIL-18 binding to hIL-18BP_ΔN-EH_ is either displayed in cartoon representation (purple, two right panels) or shown as a black dashed line representing the patch it covers on hIL-18BP_ΔN-EH_.

**Supplementary Table 1.** Molecular weights and binding parameters of recombinant hIL-18 and hIL-18BP proteins.

| Protein<br>(complex) | MW<br>(kDa) | Molecular Weight<br>determined by MALLS |  |  | Binding to hIL-18<br>determined by ITC |  |
| --- | --- | --- | --- | --- | --- | --- |
| | | Total MW<br>± SD<br>(kDa) | Protein MW<br>± SD<br>(kDa) | glycan MW<br>± SD<br>(kDa) | $K_D$<br>(95% CI)<br>(nM) | $\Delta H$<br>(95% CI)<br>(kcal/mole) |
| hIL-18 | 18.2 | 18.2 ± 0.5 | - | - | - | - |
| hIL-18BP<sub>FL</sub> | 17.6 | 32.4 ± 0.5 | 17.6 ± 0.1 | 14.8 ± 0.6 | n.d.<br>(-∞-0.245) | -19.32<br>(-18.33 - -20.32) |
| hIL-18BP<sub>ΔN</sub> | 14.4 | 24.0 ± 0.2 | 14.65 ± 0.06 | 9.3 ± 0.1 | - | - |
| hIL-18BP<sub>ΔN-EH</sub> | 14.4 | 21.12 ± 0.09 | 14.42 ± 0.02 | 6.71 ± 0.08 | n.d.<br>(-∞-0.817) | -18.13<br>(-16.91 - -19.36) |
| hIL-18<br>hIL-18BP<sub>ΔN-EH</sub> | 32.6 | 38.80 ± 0.06 | 31.9 ± 0.2 | 6.9 ± 0.2 | - | - |
| hIL-18Rα | 35.4 | - | - | - | 5.24<br>(7.94-3.38) | -26.63<br>(-25.59 - -27.77) |

MW: Molecular weight, SD: Standard Deviation, CI: Confidence interval, MALLS: Multi-Angle Laser Light Scattering, ITC: Isothermal Titration Calorimetry

